# Severity Index for Suspected Arbovirus (SISA): machine learning for accurate prediction of hospitalization in subjects suspected of arboviral infection

**DOI:** 10.1101/647206

**Authors:** Rachel Sippy, Daniel F Farrell, Daniel Lichtenstein, Ryan Nightingale, Megan Harris, Joseph Toth, Paris Hantztidiamantis, Nicholas Usher, Cinthya Cueva, Julio Barzallo Aguilar, Anthony Puthumana, Timothy Endy, Sadie J. Ryan, Anna M. Stewart Ibarra

**Affiliations:** Institute for Global Health and Translational Science, SUNY Upstate Medical University, Syracuse, NY, USA; Department of Geography, University of Florida, Gainesville, FL, USA; Department of Medicine, SUNY Upstate Medical University, Syracuse, NY, USA; Department of Microbiology, Cornell University, Ithaca, NY, USA; Teofilo Davila Hospital, Ministry of Health, Machala, El Oro Province, Ecuador

**Keywords:** machine learning, arboviral disease, dengue fever, undifferentiated febrile illness

## Abstract

**Background:** Dengue, chikungunya, and Zika are arboviruses of major global health concern. Decisions regarding the clinical management of suspected arboviral infection are challenging in resource-limited settings, particularly when deciding on patient hospitalization. The objective of this study was to determine if hospitalization of individuals with suspected arboviral infections could be predicted using subject intake data.

**Methodology/Principal Findings:** Two prediction models were developed using data from a surveillance study in Machala, a city in southern coastal Ecuador with a high burden of arboviral infections. Data was obtained from subjects who presented at sentinel medical centers with suspected arboviral infection (November 2013 to September 2017). The first prediction model—called the Severity Index for Suspected Arbovirus (SISA)— used only demographic and symptom data. The second prediction model—called the Severity Index for Suspected Arbovirus with Laboratory (SISAL)—incorporated laboratory data. These models were selected by comparing the prediction ability of seven machine learning algorithms; the area under the receiver operating characteristic curve from the prediction of a test dataset was used to select the final algorithm for each model. After eliminating those with missing data, the SISA dataset had 534 subjects, and the SISAL dataset had 98 subjects. For SISA, the best prediction algorithm was the generalized boosting model, with an AUC of 0.91. For SISAL, the best prediction algorithm was the elastic net with an AUC of 0.94. A sensitivity analysis revealed that SISA and SISAL are not directly comparable to one another.

**Conclusions/Significance:** Both SISA and SISAL were able to predict arbovirus hospitalization with a high degree of accuracy in our dataset. These algorithms will need to be tested and validated on new data from future patients. Machine learning is a powerful prediction tool and provides an excellent option for new management tools and clinical assessment of arboviral infection.

**Author Summary:** Patient triage is a critical decision for clinicians; patients with suspected arbovirus infection are difficult to diagnose as symptoms can be vague and molecular testing can be expensive or unavailable. Determining whether these patients should be hospitalized or not can be challenging, especially in resource-limited settings. Our study included data from 543 subjects with a diagnosis of suspected dengue, chikungunya, or Zika infection. Using a machine learning approach, we tested the ability of seven algorithms to predict hospitalization status based on the signs, symptoms, and laboratory data that would be available to a clinician at patient intake. Using only signs and symptoms, we were able to predict hospitalization with high accuracy (94%). Including laboratory data also resulted in highly accurate prediction of hospitalization (92%). This tool should be test in future studies with new subject data. Upon further development, we envision a simple mobile application to aid in the decision-making process for clinicians in areas with limited resources.

## Introduction

Undifferentiated febrile illness is a common clinical scenario in tropical medicine, with a long list of potential pathogens sharing similar symptoms. Arthropod-borne viruses (arboviruses), including dengue virus (DENV), chikungunya virus (CHIKV), and Zika virus (ZIKV), share common mosquito vectors (*Aedes aegypti* and *Ae. albopictus*) and often present with fever, rash, myalgias, and arthralgias. Dengue virus is endemic in the tropical Americas, and the emergence of CHIKV in 2013 in Saint Martin and ZIKV in 2015 in Brazil has brought these arboviruses to the forefront of international attention [1–3]. There were over 400,000 cases of dengue fever in Andean Latin America in 2013 [4], with transmission risk expected to increase sharply over the next 50 years [5]. Ecuador in particular has a high burden of arboviral illness, with 86,306 total cases of dengue from 2014—2018 [6–8]. There is also a high prevalence of asymptomatic DENV infections and co-infections in southern coastal Ecuador [1,9]. In 2014, CHIKV was introduced, with 35,555 cases from 2014—2018, followed by the introduction of ZIKV in 2016, with 5,304 cases from 2016—2018 [6–8].

Clinical decision-making in the context of arboviral infection is particularly challenging in resource-limited settings such as Ecuador. For instance, there may be limited healthcare professionals relative to the high disease burden, which may impact the ability to provide optimal subject care. Ecuador has 22 physicians for every 10,000 people, though this ranges by province, from 13 to 32 physicians per 10,000 people [10]. This is just above the 1 physician per 1,000 people benchmark of the World Health Organization [11], and physicians are likely concentrated in urban areas. Moreover, molecular diagnostics are often unavailable outside of large urban centers. Of Ecuador’s 4,168 healthcare establishments, 1,045 (25.1%) have a clinical laboratory [10], leaving many healthcare providers in Ecuador without these crucial diagnostic tools (*i.e.* PCR or ELISA). These infrastructural limitations create a challenging clinical environment, especially as healthcare providers need to determine whether a patient with suspected arboviral illness should be hospitalized or not. Efficient and effective triaging is essential for good clinical care in resource-limited settings [12].

Current practice in Ecuador is to hospitalize subjects suspected of dengue infection when they exhibit any of the World Health Organization (WHO) 2009 dengue warning signs, any signs of shock, or severe thrombocytopenia [13]. While treatment for dengue is supportive, proper inpatient management of severe dengue can reduce mortality dramatically [14,15]. Dengue has a wide spectrum of clinical presentations, with the majority patients recovering following a self-limited clinical course, and a small percentage progressing to severe disease characterized by plasma leakage. In the latter cases, prompt intravenous rehydration can reduce the case fatality rate to less than 1% [14]. Similarly, management of CHIKV and ZIKV infections is largely supportive, but both infecitons may result in potentially serious complications, such as adverse neonatal effects [16–19]. Deciding whether to hospitalize a subject with a suspected DENV, CHIKV, or ZIKV infection is thus an important clinical decision. This decision has other non-clinical and indirect consequences, including the utilization of hospital resources that could otherwise be used for other patients, as well as increasing the financial cost of the case when compared to less costly outpatient care. Globally, an estimated 18% of dengue cases are admitted to the hospital, with 48% managed as outpatients and 34% not seeking medical attention [20]. The average cost to manage a case of dengue is tripled if the patient is hospitalized [14,20].

Machine learning is a tool that combines statistics with computer science to make efficient use of massive data sets [21]. It differs from traditional statistical modeling (e.g. regression models) in that there are fewer assumptions about the underlying distribution of the data and the relationships between variables. While model interpretability is often a goal of traditional statistical models, this is not important in machine learning. The only goal is to create highly accurate predictions of an outcome of interest, often using as many variables as possible [22,23]. In modeling relationships with a machine learning approach, the computer incorporates connections not obvious to human beings to successfully predict an outcome of interest. Machine learning is applicable in many fields and has been previously used in medical applications, to estimate clinical risk, guide triage or diagnose disease [21,24–26]. Clinical applications of machine learning for arboviral illnesses specifically have included analysis of patient genomes for dengue prognosis [27], scanning of patient sera for DENV [28] or Zika diagnosis [29], thermal image scanning for detection of hemodynamic shock [30], analysis of body temperature patterns for diagnosis of undifferentiated fever etiology [31], and analysis of patient data for dengue fever diagnosis [25]. No studies have yet attempted to use machine learning for prediction of hospitalization among arboviral illness or undifferentiated fever patients, although it has been used to predict critical care and hospitalization outcomes based on emergency department triage data in children and adults [32,33].

The objective of this study was to determine if the hospitalization of individuals with suspected arboviral infections could be predicted using subject intake data. In this study, we take a retrospective view of arboviral infection management in a tropical city in southern coastal Ecuador using data from an ongoing prospective surveillance study. Using actual clinical practice as a guide, we assessed the ability of seven machine learning algorithms to determine hospitalization using basic symptom and demographic data that was collected via standard intake of subjects with suspected DENV, CHIKV, or ZIKV infections. The machine learning approach and algorithms developed here could potentially support physicians faced with complex clinical management decisions in areas where multiple arboviruses co-circulate, such as Ecuador.

## Methods

### Ethics statement

This study protocol was reviewed and approved by Institutional Review Boards at the State University of New York (SUNY) Upstate Medical University, the Luis Vernaza Hospital in Guayaquil, Ecuador, and the Ecuadorean Ministry of Health (MoH). Clinical and demographic data from study subjects was obtained following informed consent (and/or assent, as applicable) per the protocol of an ongoing arboviral surveillance study in Machala, Ecuador (described previously) [1].

### Study design and data source

We conducted a retrospective analysis of data from a prospective arbovirus surveillance study, which included subjects (age >6 months) recruited from Ecuadorean MoH clinical sites from November 2013 to September 2017 in Machala, Ecuador. Subjects were identified as a part of an ongoing, multi-year arbovirus surveillance project, a description of which has been published previously [1]. Briefly, subjects were invited to enroll in the study if they presented at the reference hospital or one of four outpatient clinics and were diagnosed with arboviral infection by MoH physicians. In 2014 and 2015, we recruited subjects who were clinically diagnosed with dengue fever by MoH physicians. Following the emergence of CHIKV (2014) and ZIKV (2016), the inclusion criterion in 2016 and 2017 was expanded to include subjects clinically diagnosed with DENV, CHIKV or ZIKV infection. At the time of enrollment, subject demographic information, clinical history, and symptoms present during current illness were collected using a questionnaire administered by trained study personnel. Subjects were asked about symptoms in the past 7 days: headache, anorexia or nausea, muscle or joint pain, rash, bleeding (defined as bleeding from respiratory, digestive, or genitourinary mucosa), rhinorrhea, vomiting, lethargy or drowsiness, cough, abdominal pain, diarrhea, and retro-orbital pain. Conjunctivitis was later added to the enrollment survey after the emergence of ZIKV but was not included in this analysis. Laboratory data (hematocrit, white blood cell count, neutrophils, lymphocytes, and platelet count) was collected at the time of enrollment if the subject had copies of recent laboratory evaluation (for outpatients), or the first labs on admission to the hospital (for hospitalized subjects) were used. Additional laboratory data were available for hospitalized subjects, but analysis was limited to the aforementioned three parameters as these were consistently available among a subset of the non-hospitalized subjects. Laboratory arboviral diagnostic data were not included, as these data are often not available at the time that a physician decides whether or not to hospitalize a patient, and we utilized only rely the data that would realistically be available. Data from enrollment surveys was used for the analysis of the non-hospitalized outpatients in the current study. Laboratory data on the hospitalized subjects was verified by review of medical records and managed using REDCap software [34] hosted at SUNY Upstate Medical University.

### Exclusion criteria

Hospitalized subjects whose physical medical records could not be located and subjects with incomplete enrollment survey data (*i.e.* missing hospitalization status, symptom survey questions) were excluded. The subset of the non-hospitalized subjects who had available laboratory data were included in a second analysis with the same hospitalized cohort, all of whom had available laboratory data.

### Statistical analysis

The outcome variable was hospitalization status. Variables of interest included demographic data, presenting symptoms, past medical history, and laboratory data (hematocrit, white blood cell count, neutrophils, lymphocytes, and platelet count). A prediction algorithm was developed using demographic and symptom data only (28 total predictors), called the Severity Index for Suspected Arbovirus (SISA, in Spanish the *Severidad de Infecciones Sospechosas por Arbovirus*). A second prediction algorithm was developed using demographic, symptom, and laboratory data (33 total predictors), called the Severity Index for Suspected Arbovirus with Laboratory (SISAL, in Spanish the *Severidad de Infecciones Sospechosas por Arbovirus con datos del Laboratorio*). Characteristics for hospitalized and non-hospitalized subjects among these subject groups were compared using a two-sample t-test (continuous) or Fisher’s exact test (categorical).

In machine learning, 10-fold cross-validation with holdout data results in an unbiased estimate of model validity and accuracy [22,35]. Thus, our datasets were divided by random sampling into training and testing (holdout) data sets. For SISA, the training set was 85% of the full dataset and the testing set was the remaining 15%. For SISAL, the training set was 70% of the full dataset and the testing set was the remaining 30%. With the training dataset, we used repeated 10-fold cross validation to test the ability of six algorithms with diverse statistical approaches— bagged trees (bags) [36], *k* nearest neighbors regression (knn) [37], random forest [38], elastic net regression [39], generalized boosting models (gbm) [40], and neural networks [41]—to predict hospitalization. Because we have no prior assumptions about the nature of the relationship between the available predictors and the outcome, we use a variety statistical approaches to improve the likelihood that we will find an algorithm that works well with these data. Following a published criticism of machine learning prediction compared to logistic regression [42], we added logistic regression to our list of algorithms to test (seven total algorithms). For models with tuning parameters (knn, random forest, elastic net, and gbm), tuning was performed using another layer of 10-fold cross-validation [43]. Each algorithm was then used to predict hospitalization outcomes within the testing dataset. Measures of discrimination [44], including accuracy, Cohen’s kappa, and area under the curve (AUC) for the receiver operating characteristic (ROC) were calculated for each of the ten cross-validations and summarized as mean values. The best algorithm for each dataset was chosen based on the highest AUC as calculated from the test set. Model residual plots were examined. Data analysis and visualization was performed using SAS version 9.2 (SAS Institute, Cary, NC) and R version 3.2.2 (R Foundation for Statistical Computing, Vienna, Austria) including packages haven [45], caret [37], MASS [41], ipred [36], randomForest [38], elasticnet [39], gbm [40], nnet [41], mgcv [46,47], kernlab [48], glmnet [49], and pROC [50].

We compared the prediction abilities of SISA versus SISAL to assess whether laboratory data could improve our ability to predict subject hospitalization status. Because there may be some selection bias for subjects with available laboratory data (e.g. more severe symptoms, more similar subject data, or different socioeconomic status compared to typical patients with clinical arboviral diagnosis), the subject groups in SISA and SISAL may not be exchangeable [51]. We performed a sensitivity analysis to determine if the selected algorithm and prediction ability of SISA is the same when using all SISA subjects or SISAL subjects (without laboratory data) for the training and testing steps.

## Results

### General Characteristics

Between November 20, 2013 to September 13, 2017, 592 subjects were recruited into the arboviral surveillance study. After exclusions (Figure 1), 534 subjects were included in the dataset for SISA, of which 59 were hospitalized and 475 were not hospitalized. The SISA training dataset included 455 subjects and the test dataset included 79 subjects. The SISAL dataset included 98 subjects, of which 59 were hospitalized and 39 were outpatients. The SISAL training dataset included 70 subjects and the test dataset included 28 subjects. Demographics and symptoms for the two datasets are in Table 1. Presenting temperature was higher, and presence of mucosal bleeding, vomiting, and abdominal pain were significantly more common in hospitalized subjects in the SISA dataset.

**Figure 1:**
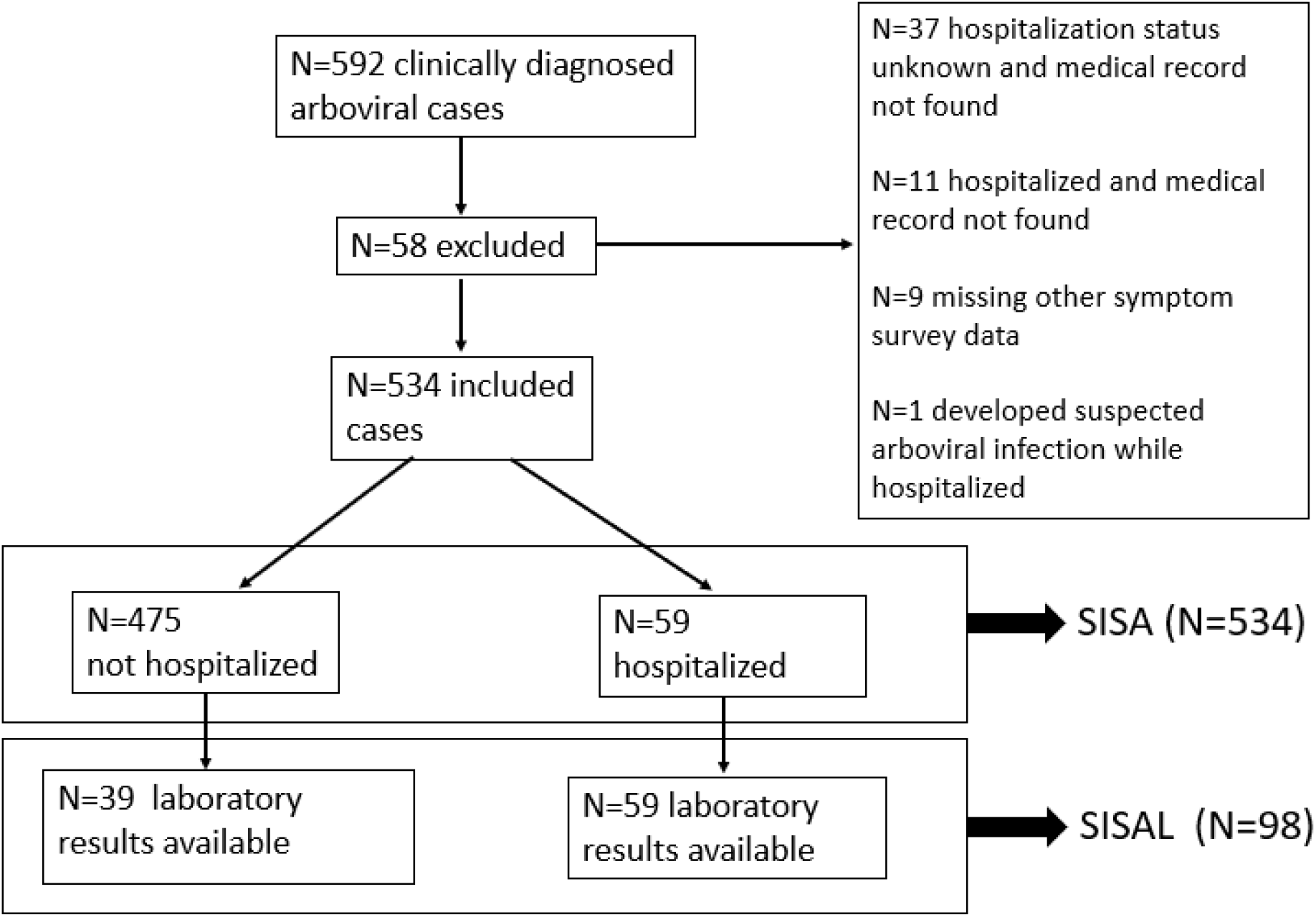
Flow Diagram of Subject Selection. Subjects clinically diagnosed with arboviral (dengue, chikungunya, Zika) infections were recruited from sentinel clinics in Machala, Ecuador. A subset of subjects were used to test the Severity Index for Suspected Arbovirus (SISA) and Severity Index for Suspected Arbovirus with Laboratory (SISAL) machine learning algorithms.

**Table 1.** Overview of demographics, symptoms, and laboratory values for subjects enrolled in the study (n=543). Numerical data are shown as means and were analyzed with Welch 2-Sample T-test. Categorical data shown as percentages and analyzed with Fisher’s Exact Test. Bold p-values indicate p<0.05.

| SISA (N=543) |  |  |  | SISAL (N=98) |  |  |
| --- | --- | --- | --- | --- | --- | --- |
|  | Hospitalized<br>(n=59) | Outpatient<br>(n=484) | p-value | Hospitalized<br>(n=59) | Outpatient<br>(n=39) | p-value |
| Age (years) | 23.3 | 25 | 0.38 | 23.3 | 22.3 | 0.75 |
| Height (cm) | 154.2 | 147.9 | 0.01 | 154.2 | 154.2 | 0.99 |
| Weight (kg) | 58.2<sup>A</sup> | 56.5 | 0.61 | 58.2<sup>A</sup> | 57 | 0.78 |
| MUA Circumference<br>(cm) | 25.4 | 25.9 | 0.5 | 25.4 | 25.9 | 0.64 |
| Waist Circumference<br>(cm) | 83.9 | 78.4 | 0.05 | 83.9 | 78 | 0.1 |
| Temperature (°C) | 37.7 | 37.4 | 0.01 | 37.7 | 37.2 | 0.02 |
| Hematocrit (%) |  |  |  | 37.6 | 39.8 | 0.02 |
| WBC Count (cells/μL) |  |  |  | 5435 | 6863 | 0.18 |
| Neutrophils (%) |  |  |  | 58.1 | 54.5 | 0.38 |
| Lymphocytes (%) |  |  |  | 30.6 | 33.5 | 0.38 |
| Platelet Count |  |  |  | 117627 | 203333 | <0.01 |
| Gender (% female) | 63 | 54 | 0.27 | 63 | 51 | 0.30 |
| Fever in past 7 days | 97 | 94<sup>C</sup> | 0.56 | 97 | 97 | 1.00 |
| Head pain | 64 | 79<sup>A</sup> | 0.01 | 64 | 82 | 0.07 |
| Nausea | 54 | 53<sup>A</sup> | 0.09 | 54 | 59 | 0.68 |
| Muscle or joint pain | 70 | 81<sup>A</sup> | 0.04 | 70 | 82 | 0.24 |
| Rash | 20 | 29<sup>A</sup> | 0.22 | 20 | 15 | 0.60 |
| Bleeding | 24 | 4<sup>A</sup> | <0.01 | 24 | 5 | 0.02 |
| Rhinorrhea | 19 | 26<sup>A</sup> | 0.27 | 19 | 21 | 1.00 |
| Vomiting | 56 | 27<sup>B</sup> | <0.01 | 56 | 31 | 0.02 |
| Drowsiness or lethargy | 80 | 84<sup>B</sup> | 0.46 | 80 | 92 | 0.15 |
| Coughing | 37 | 35<sup>B</sup> | 0.77 | 37 | 26 | 0.27 |
| Abdominal pain | 70 | 50<sup>B</sup> | <0.01 | 70 | 56 | 0.20 |
| Diarrhea | 29 | 22<sup>B</sup> | 0.33 | 29 | 44 | 0.19 |
| Retroorbital pain | 56 | 66<sup>C</sup> | 0.15 | 56 | 77 | 0.05 |
| Positive tourniquet test | 17 | 3<sup>D</sup> | <0.01 | 17 | 5 | 0.01 |
| History of allergies | 20 | 20<sup>A</sup> | 0.86 | 20 | 18 | 0.80 |
| History of hypertension | 5 | 6<sup>A</sup> | 1.00 | 5 | 10 | 0.43 |
| History of asthma | 2 | 4<sup>A</sup> | 0.71 | 2 | 3 | 1.00 |
| History of cancer | 0 | 2<sup>A</sup> | 0.61 | 0 | 0 | 1.00 |
| History of diabetes | 2 | 3<sup>A</sup> | 1.00 | 2 | 0 | 1.00 |
| History of dengue in<br>household | 17 | 12<sup>A</sup> | 0.30 | 17 | 5 | 0.12 |
| History of dengue | 19 | 22<sup>A</sup> | 0.74 | 19 | 18 | 1.00 |
| Pregnancy (self-<br>reported) | 19 | 2 | <0.01 | 19 | 3 | 0.22 |

SISA= Severity Index for Suspected Arbovirus or Severidad de Infeccion Sospechosa de Arbovirus, SISAL= Severity Index for Suspected Arbovirus with Laboratory or Severidad de Infeccion Sospechosa de Arbovirus con Laboratorio, cm=centimeters, kg=kilograms, MUA=mid- upper arm, °C=degrees Celsius, WBC=white blood cell, μL=microliters. <sup>A</sup> Missing n=1, <sup>B</sup> Missing n=2, <sup>C</sup> Missing n=3, <sup>D</sup> Missing n=4, % Pregnant is taken from the total population (male and female)

### Prediction of Hospitalization Status

Accuracy, Cohen’s kappa, and AUC for the training and test sets are shown in Figure 2. For SISA, using only symptoms and demographics, generalized boosting model, elastic net, neural networks, and logistic regression performed well with the test set (accuracy: 89.8—96.2%, Cohen’s kappa: 0.00—0.77, AUC: 0.50—0.91). The generalized boosting model algorithm was found to have the best AUC (0.91) among the test dataset and was the second-best algorithm in the training set. The sensitivity for this algorithm was 95.8%, and the specificity was 87.5% when predicting hospitalization of subjects in the test dataset. The calibration plot for this prediction is in Figure 3. The variables with the greatest influence on the final SISA model were drowsiness, bleeding, and temperature.

**Figure 2:**
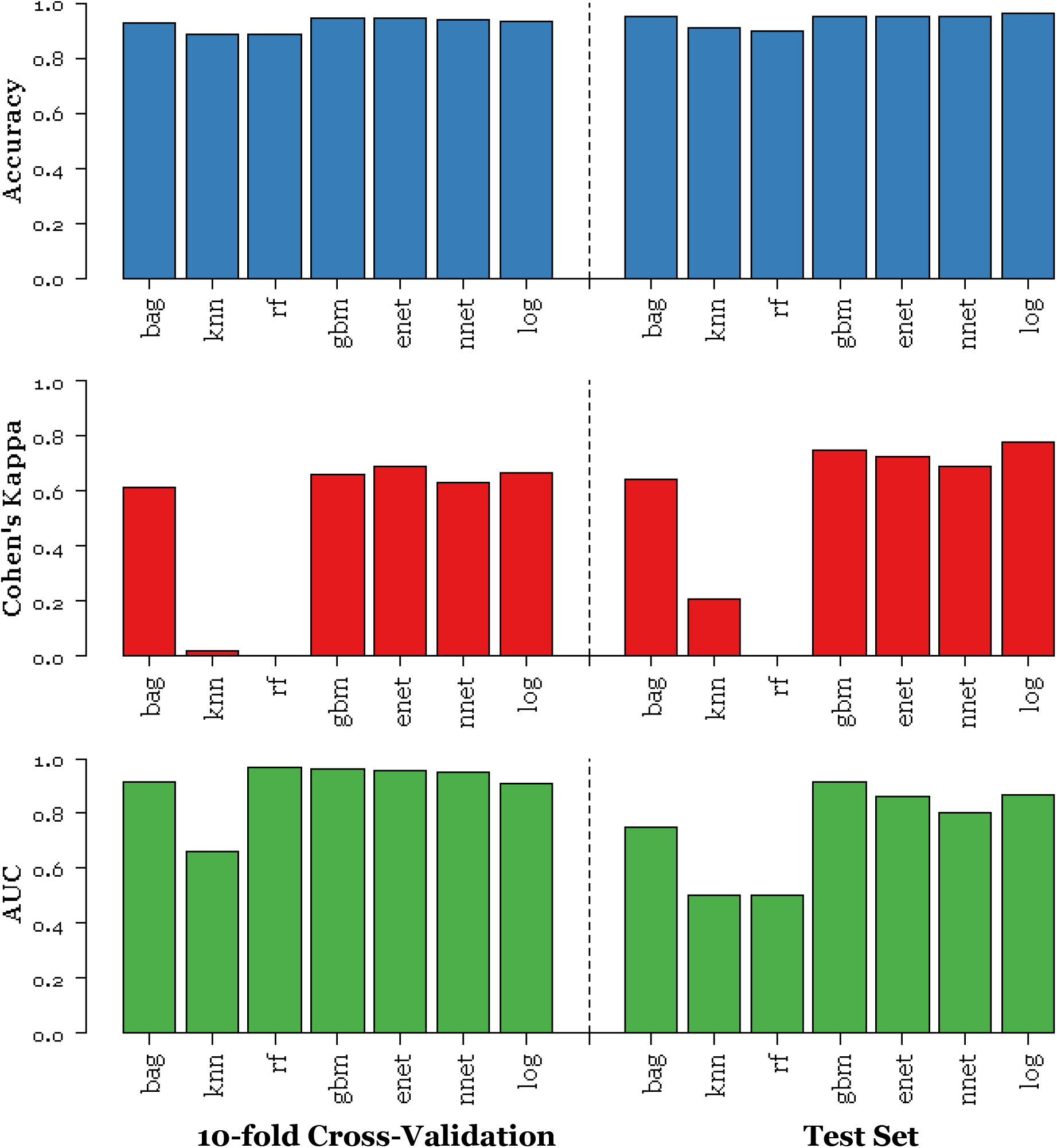
Results for SISA Dataset. Accuracy (blue), Cohen’s kappa (red), and AUC (green) were calculated for the 10-fold cross validation (left) and the holdout test dataset (right) for prediction of hospitalization status in clinically diagnosed dengue, chikungunya or Zika virus infections. bag=bagged trees, knn=*k* nearest neighbors, rf=random forest, gbm=generalized boosting models, enet=elastic net, nnet=neural networks, log=logistic regression

**Figure 3:**
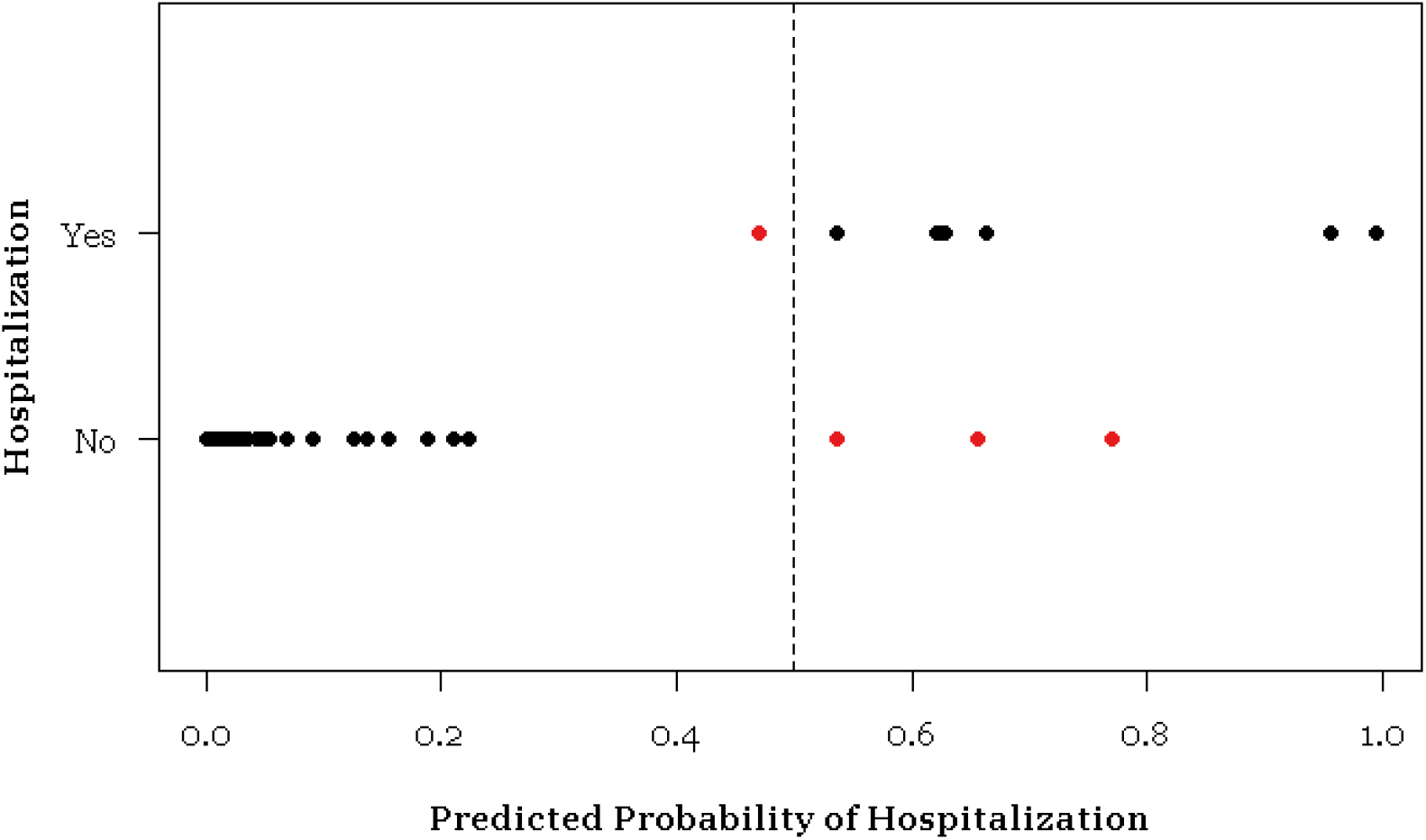
Calibration Plot for SISA Prediction. For the final SISA algorithm (generalized boosting model), the model prediction probability is compared to the actual hospitalization outcome among the test set. Incorrect predictions are in red: there were three subjects incorrectly classified for hospitalization and one subject incorrectly classified for non-hospitalization.

Results for SISAL, where laboratory parameters were included as well as symptoms and demographics, are in Figure 4. All models except neural networks and *k* nearest neighbors performed well with the test set (accuracy: 64.3—92.6%, Cohen’s kappa: 0.25—0.85, AUC: 0.62—0.94). The elastic net algorithm had the best AUC (0.94) among the test dataset and was the third-best algorithm in the training set. The sensitivity for SISAL was 100% and the specificity was 88.2% when predicting hospitalization of subjects in the test dataset. The calibration plot for this prediction is in Figure 5. The variables with the greatest influence on the final SISAL model were drowsiness, orbital pain, and platelet count.

**Figure 4:**
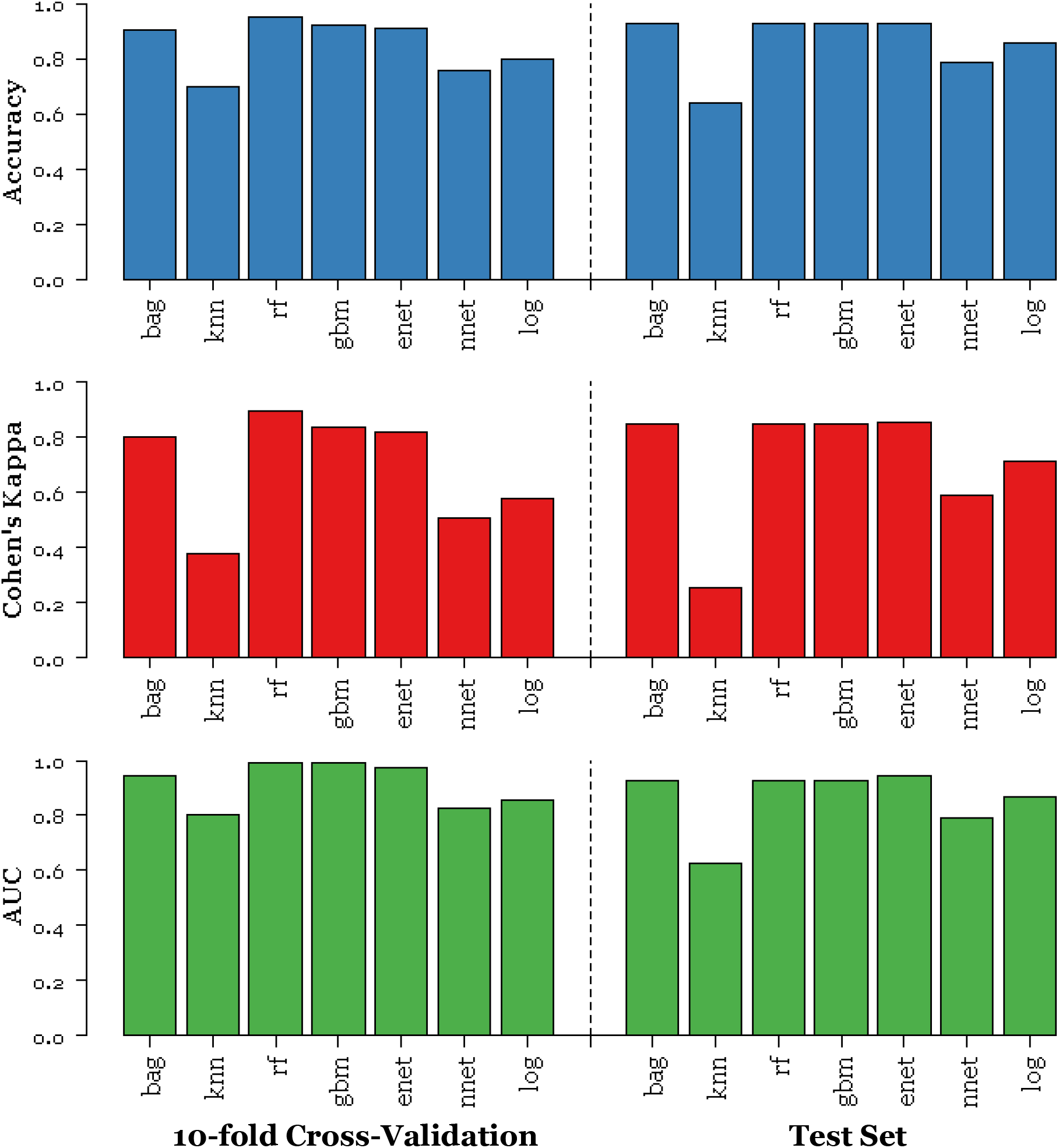
Results for SISAL Dataset. Accuracy (blue), Cohen’s kappa (red), and AUC (green) were calculated for the 10-fold cross validation (left) and the holdout test dataset (right) for prediction of hospitalization status in clinically diagnosed dengue, chikungunya or Zika virus infections. bag=bagged trees, knn=*k* nearest neighbors, rf=random forest, gbm=generalized boosting models, enet=elastic net, nnet=neural networks, log=logistic regression

**Figure 5:**
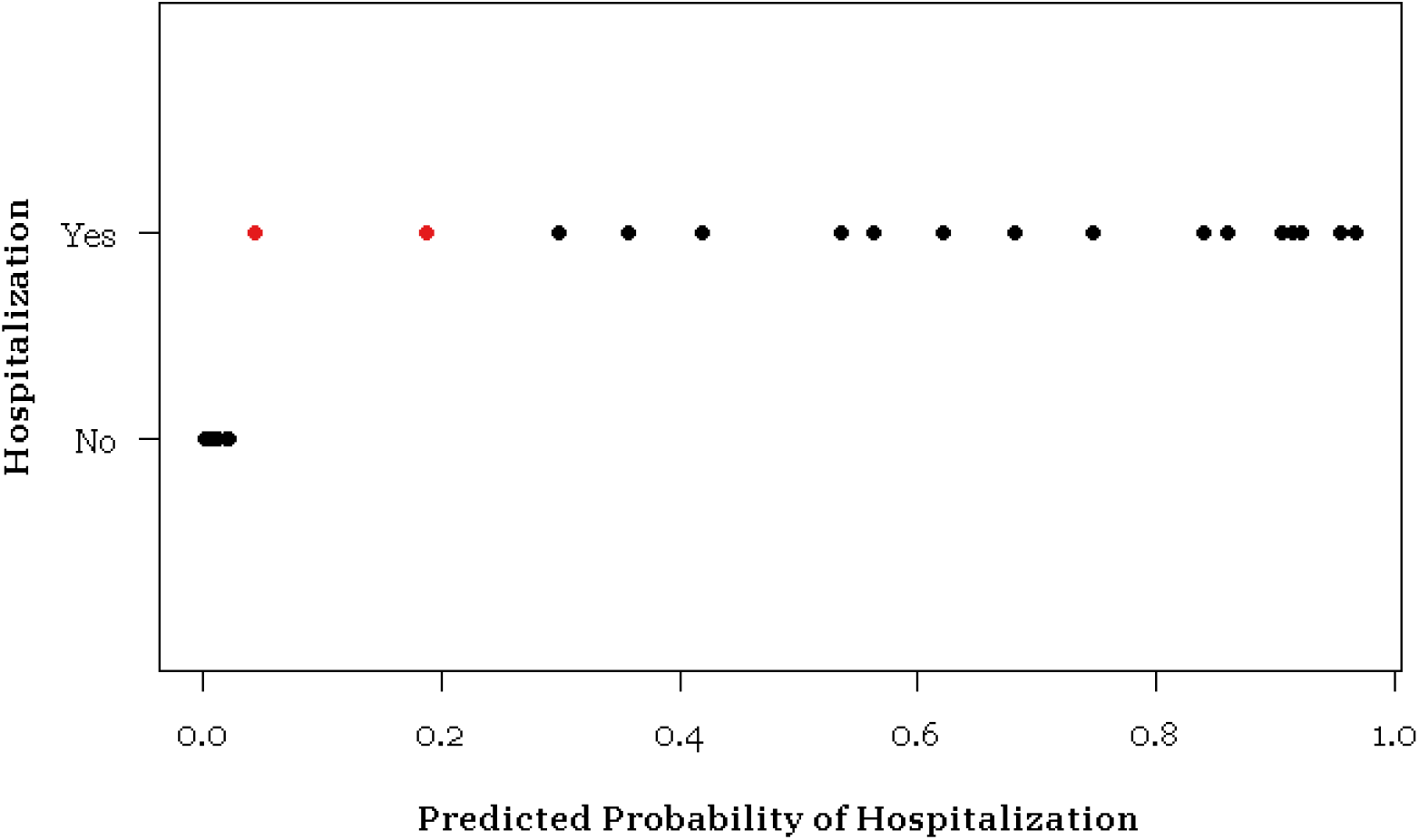
Calibration Plot for SISAL Prediction. For the final SISAL algorithm (elastic net regression), the model prediction probability is compared to the actual hospitalization outcome among the test set. Incorrect predictions are in red: there were two subjects incorrectly classified for hospitalization.

The results for SISA when trained with the SISAL subjects (without laboratory data) are in the S1 Figure. All models except *k* nearest neighbors and logistic regression performed well (test set accuracy: 53.5—92.9%, Cohen’s kappa: 0.04—0.86, AUC: 0.51—0.94). The bagged trees, random forest, generalized boosting models, and elastic net algorithm had identical AUC (0.94). The sensitivity was 100% and the specificity was 88.2% when predicting hospitalization of subjects in the test dataset. If the SISA and SISAL subjects were exchangeable, we would expect the SISAL subject group (without laboratory data) to produce the same results as the SISA analysis. Because these results differ from those obtained in the SISA analysis, we conclude that the SISAL subjects are not exchangeable with the SISA subjects.

## Discussion

Suspected arboviral infections impose large health and financial burdens on populations in which the diseases are endemic. In 2013, the estimated global cost of dengue illness was US $8—9 billion [20]. In many arbovirus endemic regions, DENV, CHIK, and ZIKV infections are diagnosed based on clinical presentation and basic laboratory results, which can be difficult due to nonspecific symptoms and limited availability of definitive diagnostic tools [52]. In this study, we demonstrate that our machine learning algorithms were able to predict hospitalization status among our cohort of subjects with suspected arboviral illness with up to 96% accuracy using only symptom and demographic data. We thus describe the early development of a new tool, SISA/SISAL, which in the future may be utilized by clinicians in resource-limited settings when triaging subjects with suspected arboviral illness.

In our cohort, we found that hospitalized cases had statistically significant - though clinically insignificant - elevations in temperature at presentation in both SISA and SISAL. This demonstrates an algorithm’s ability to make use of data that might be overlooked by the clinician. In the SISA dataset, mucosal bleeding, vomiting, and abdominal pain were more common in hospitalized subjects than in outpatients. In the SISAL dataset, while hospitalized subjects experienced more mucosal bleeding and vomiting than outpatients, presence of abdominal pain did not differ between groups. This could suggest that the outpatient subjects who were sent for laboratory testing represented cases of serious concern, as abdominal pain may qualify those with suspected or confirmed dengue for hospital admission [13,14]. For SISAL, hospitalized subjects had lower hematocrit and platelet counts when compared to non-hospitalized subjects, with the lower platelet counts to be expected in hospitalized dengue cases. The WBC counts showed no significant difference, which could be due to leukocytosis or leukopenia obscuring the mean difference between hospitalized and non-hospitalized subjects.

Our sensitivity analysis revealed that SISA produced different results when its training/testing dataset was restricted to those subjects with laboratory data available. This result is unsurprising, as we expect that selection bias is contributing to the subjects available for the SISA and SISAL datasets. All hospitalized subjects had laboratory data available, and we would additionally expect that subjects with laboratory data had some signs or symptoms that would prompt the attending physician to request laboratory diagnostics, setting them apart from subjects without laboratory data. These signs and symptoms are also likely linked to whether subjects were eventually hospitalized or not, meaning these groups of subjects are not directly comparable to one another. When we used the SISA approach (symptoms and demographics only) for a dataset comprised of the SISAL group of subjects (without laboratory data), we found that the AUC was identical to the AUC from the SISAL approach. This would suggest that we are unable to improve our prediction of hospitalization status by using subject laboratory data. The AUC was higher for this group of subjects, but these improvements are likely due to fundamental differences between the SISA/SISAL groups of subjects. These patient groups should continue to be analyzed with separate algorithms.

This is the first use of machine learning to predict hospitalization status of subjects with clinically diagnosed arboviral infections. Our models exhibit high accuracy, sensitivity, and specificity in a region with a high burden of co-circulating of DENV, CHIKV, and ZIKV. These algorithms, particularly SISA, use information that could easily be obtained in resource-limited settings, suggesting the potential to develop a useful tool for clinicians. Our model’s accuracy is consistent with tools previously reported in the literature. Past predictive modeling of disease with a machine learning approach had been efficacious in the diagnosis of pneumonia (95% sensitivity, 96.5% specificity), dengue (70% sensitivity, 80% specificity), hepatitis (96% accuracy), and tuberculosis (95% accuracy) using clinical and laboratory parameters [25,53–55].

There has been criticism regarding the use of machine learning in prediction models: a recent systematic review found that machine learning predictions had no advantage over logistic regression predictions on average [42]. Christodoulou et al. do an excellent job of outlining some common missteps in the use of machine learning for prediction and the somewhat alarming lack of transparency in many published machine learning prediction models. We agree with many of the assertions made by the authors and strive to improve reporting and validation in our own work as a result. However, in this specific study, we did not find that logistic regression performed better than other algorithms. Our overall approach differs from that of most machine learning papers in that we did not assume that one particular algorithm would have superior prediction abilities for our data, but rigorously compared multiple algorithms with the goal of finding an algorithm that functions well with our predictors and outcome of interest, to be further validated with a new dataset in future research. We have no illusions about the potential lack of generalizability of our data and caution against any strong conclusions about the future utility SISA/SISAL in predicting hospitalization status for future patients. We present the current study as preliminary yet promising results in the development of a future tool that will need additional, vigorous validation in future sets of subject data before use in the real world.

Numerous studies have looked at clinical and laboratory findings specific to certain arbovirus diagnoses, yet few have proposed tools that can aid in management of unconfirmed febrile illness [56–59]. A study in Puerto Rico of acute febrile illness emergency room cases found the tourniquet test and leukopenia to be predictive of dengue diagnosis, yet dengue was confirmed in only 11% of their total 284 cases [60]. In Thailand, fever, positive tourniquet test and leukopenia differentiated confirmed dengue from other febrile illness initially suspected as dengue [61]. Also in Thailand, among a sample of 172 children with acute fever without obvious cause, those with dengue had several laboratory parameters that differentiated them from the other febrile illness [52]. While these studies were able to distinguish dengue from other acute febrile illness, they highlight the large proportion of cases that do not get a confirmed diagnosis and have not moved beyond initial reports to demonstrate predictive abilities. With SISA/SISAL, the approach is more empirical. Clinical diagnosis of DENV, CHIKV, or ZIKV was a starting point for the machine learning used here. Given that timely laboratory diagnostics may not be available, grouping these suspected subjects reflects the reality that physicians face in the clinic in arbovirus-endemic regions. That such a model can accurately predict hospitalization outcome suggests that SISA/SISAL could be expanded to undifferentiated fever. The ability of machine learning models to predict hospital admission outcomes using only emergency department triage data lends support to expanding our approach to undifferentiated fever [32,33]. Of the suspected arboviral cases analyzed here, approximately 54% were confirmed as acute or recent DENV infection, 17% had acute CHIKV infection, and 29% had no pathogen identified (based on analysis of subjects in 2014 and 2015) [1]. Results of the 2016 and 2017 subject samples are pending, but preliminary PCR testing suggests predominance of CHIKV in 2016 and ZIKV in 2017.

Clinicians rely on tools to help make decisions about patient management, and simple tools are can benefit physicians in limited-resource settings [62,63]. Smart phones are commonly used in Ecuador and mobile health tools are a great option for physicians, with several popular apps that include various triage rules and scores, such as MDCalc [64]. After further development and validation of our algorithmic approach, and evaluation of its potential benefit in the clinic, we conceive of its inclusion in a user-friendly mobile application to aid in the decision to hospitalize patients with undifferentiated fever.

### Limitations

The variables with the greatest influence on the final SISA model were drowsiness, temperature, and nausea; for the SISAL model they were drowsiness, orbital pain, and platelet count. An important caveat inherent to the nature of machine learning is that the exact weight of each variable in the final prediction model is difficult to assess and interpret, thus we cannot propose a causal relationship or correlation between these variables and our outcome of hospitalization.

The SISA/SISAL models are presented here in the first iteration of their use. They have not yet been validated beyond the current datasets, but the use of holdout data and 10-fold cross-validation provides us with an unbiased estimate of model validity as well as prediction accuracy. An external validation of these algorithms with a new dataset is ongoing, as well as the testing of fewer prediction variables with the eventual goal of an easy-to-use online or mobile app for use in the clinic.

In this study, we used the outcome of subject hospitalization for both prediction models. The sensitivity and specificity of SISA/SISAL relies on the assumption that the subjects in this dataset were correctly hospitalized. It is possible that some subjects were treated as outpatients when they should have been hospitalized, or that some subjects were hospitalized unnecessarily. For subjects that were incorrectly treated as outpatients, it is unlikely that the subject would not have returned to a clinic to receive care, as their symptoms would likely drive them to do so. Because our collection of medical records was retrospective, we were able to capture subject hospitalization at any point, even if they were initially treated as outpatients. Hospital Teofilo Davila is the reference MoH hospital in the province, and it is unlikely that these subjects would have sought care at a hospital elsewhere. It is possible for some subjects to have been hospitalized unnecessarily; we have no way of identifying these subjects or truly knowing if it was safe for these subjects to have been treated as outpatients. As a result, our algorithms could thus recommend hospitalization unnecessarily. Although hospitalization could place undue financial burden on some patients and the health system, failure to hospitalize a serious case could results in grave consequences and we would prefer to take a cautious approach in hospitalization decision-making. Moreover, these algorithms are merely intended as a tool to inform clinical judgement, not to replace important clinical triage decisions [65].

The time period during which our data were collected (2014—2017) included the emergence of two important new arboviruse—CHIKV and ZIKV. The potential severity of these infections and their novelty may have increased the number of patients willing to be hospitalized or to seek healthcare in the first place. With ZIKV infection, physicians may have been more likely to hospitalize pregnant women. This may limit the generalizability of SISA/SISAL in future subject datasets, though as viral diseases continue to emerge globally, it is important to test the ability of decision-making tools to function under these dynamic scenarios. For new diseases with clear warning signs for potentially severe disease, we would expect SISA/SISAL to work well.

## Conclusions

Clinicians in resource-limited settings commonly encounter subjects with a suspected diagnosis of DENV, CHIKV, or ZIKV infection and often have limited tools at their disposal. A subject may be unable or unwilling to provide a laboratory specimen, and diagnostic testing may not always be available. The SISA/SISAL models are promising clinical tools, given the high sensitivity and specificity for both models. Machine learning, if used thoughtfully, can be a powerful method for building such prediction models, making the best use of real-world available clinical data.

## Acknowledgements

Many thanks to the Ministry of Health of Ecuador and SUNY Upstate’s Institute for Global Health and Translational Science, especially Tina Lupone, as well as the Upstate team in Machala.

## Supporting information captions

**S1 Figure: SISA Analysis of SISAL Dataset** Accuracy (blue), Cohen’s kappa (red), and AUC (green) were calculated for the 10-fold cross validation (left) and the holdout test dataset (right) for prediction of hospitalization status in clinically diagnosed DENV, CHIKV or ZIKV infections. bag=bagged trees, knn=*k* nearest neighbors, rf=random forest, gbm=generalized boosting models, enet=elastic net, nnet=neural networks, log=logistic regression, DENV=dengue virus, CHIKV=chikungunya virus, ZIKV=Zika virus

