## Supplementary figures and images for "Severity Index for Suspected Arbovirus (SISA): machine learning for accurate prediction of hospitalization in subjects suspected of arboviral infection" (from another version of this preprint: version 1, 2019-05-24)

### S1 Figure

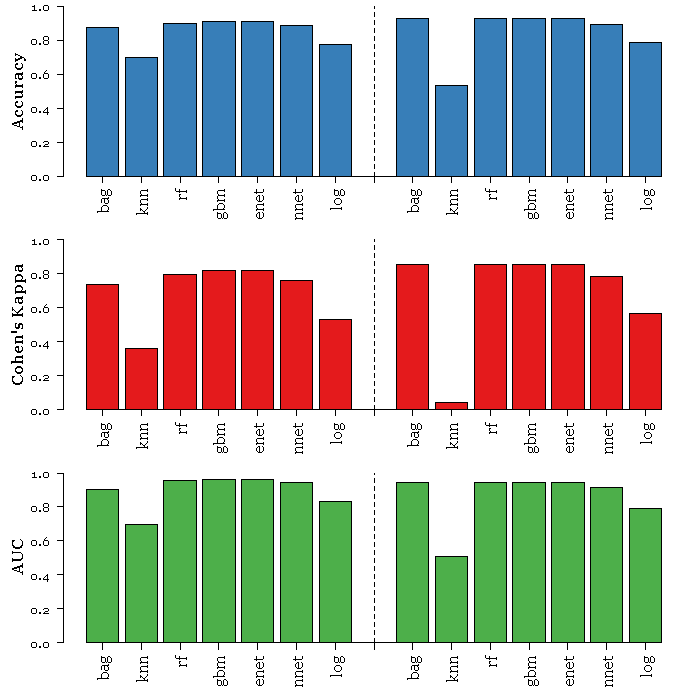

**10-fold Cross-Validation	Test Set**
